# Paralog-specific intrabodies for PSD-93 and SAP102 expand the molecular toolkit to resolve excitatory synapse organization

**DOI:** 10.1101/2025.08.24.671450

**Authors:** Christelle Breillat, Ellyn Renou, Manon Darribere, Charlotte Rimbault, Vincent Talenton, Agathe Ecoutin, Sophie Daburon, Christel Poujol, Daniel Choquet, Cameron Mackereth, Matthieu Sainlos

## Abstract

A scarcity of live, paralog-specific tools has limited analysis of PSD-MAGUKs at excitatory synapses. To address this gap, we engineered small, ^10^FN3-derived binders that selectively recognize PSD-93 and SAP102 -alongside an enhanced PSD-95 reagent- and converted them into regulated, gene-encoded intrabodies for endogenous imaging. Through sequence-guided selection and targeted optimization, we obtained high-specificity reagents that label their native targets in neurons with minimal perturbation and support multiplexed live-cell and advanced imaging modalities. This toolkit enables differential visualization of MAGUK paralogs at native levels and provides a practical route to dissect their distinct contributions to synapse organization and plasticity.

## Introduction

Excitatory synapses are molecularly dense, heterogeneous assemblies whose nanoscale organization and plasticity emerge from dynamic protein–protein interaction (PPI) networks.(**Bayes *et al***., **2011; Dieterich and Kreutz, 2016; Sudhof and Malenka, 2008**) Within this complexity, postsynaptic density (PSD) scaffolds of the MAGUK/DLG family—PSD-95 (DLG4), PSD-93 (DLG2), SAP102 (DLG3) and SAP97 (DLG1)—play central organizational roles(**Cizeron *et al***., **2020; Won *et al***., **2017**) but are challenging to dissect individually because of their high sequence homology and conserved PPI modules. As a result, the discrete functions of each paralog remain difficult to resolve with conventional perturbations or labels.

Protein-engineered binders provide an alternative route.(**Choquet *et al*.,2021; D’Este *et al*.,2024; Muyldermans, 2021; Trimmer, 2022**) We previously evolved ^10^FN3-based monobodies(**Koide and Koide, 2007; Sha *et al*.,2017**) that bind the PSD-95 PDZ1–2 supramodule with sub- to micromolar affinities, recognize epitopes outside the canonical PDZ binding groove, and show strong selectivity over other MAGUK paralogs in cells.(**Rimbault *et al*.,2019**) In particular, single-residue differences such as PSD-95 F119 (R in other paralogs) were identified as key specificity determinants. Building on this, we recently established minimally interfering, gene-encoded PSD-95 probes by coupling the binders to a transcriptional repression/zinc-finger regulation cassette that matches probe expression to target abundance, enabling robust imaging across modalities (STED, PALM, DNA-PAINT) without detectable perturbation of synaptic physiology.(**Rimbault *et al*.,2024**)

Here, we generalize this strategy to other PSD-95 paralogs. Guided by sequence analysis of the PDZ1–2 supramodule and the distribution of paralog-unique residues, we redesigned ^10^FN3 libraries and introduced a depletion step against non-target paralogs to enforce specificity during phage selections. This yielded lead binders for PSD-93 and SAP102, alongside a high-affinity PSD-95 clone. We then validated full-length target engagement in cells using FRET/FLIM and converted these binders into regulated intrabody probes for imaging endogenous proteins in neurons.

## Results

### Targets, libraries and selection strategy design

In order to complement our PSD-95 specific binders, Xph15 and Xph20, with binders that recognize the other paralogs, PSD-93, SAP102 and SAP97, we first considered the same targeted domains used for PSD-95 (**Fig. 1a**).(**Rimbault *et al*.,2019**) Indeed, as noted previously, the first two PDZ domains of the PSD-95 MAGUK family present the advantage of having a highly conserved functional region, *i*.*e*., the PDZ domain binding groove, which is the center of this PPI module activity as it is where it engages with PDZ domain partners. In consequence, isolation of paralog-specific binders, as demonstrated for Xph15 and Xph20,(**Rimbault *et al*.,2024; Rimbault *et al*.,2019**) is associated with epitopes distant from the binding groove and therefore their interactions are not impairing the PDZ domain PPI function. Analysis of the paralogs tandem domains sequences clearly indicated the presence of unique residues that could be used as recognition handles for the generation of specific binders as was the case for the unique PSD-95 F119 (arginine residue in all other proteins), which was present in Xph15, Xph18 and Xph20 epitopes. We therefore produced the four tandem PDZ domains in an enzymatically biotinylated format for presentation as targets.

**Fig. 1.**
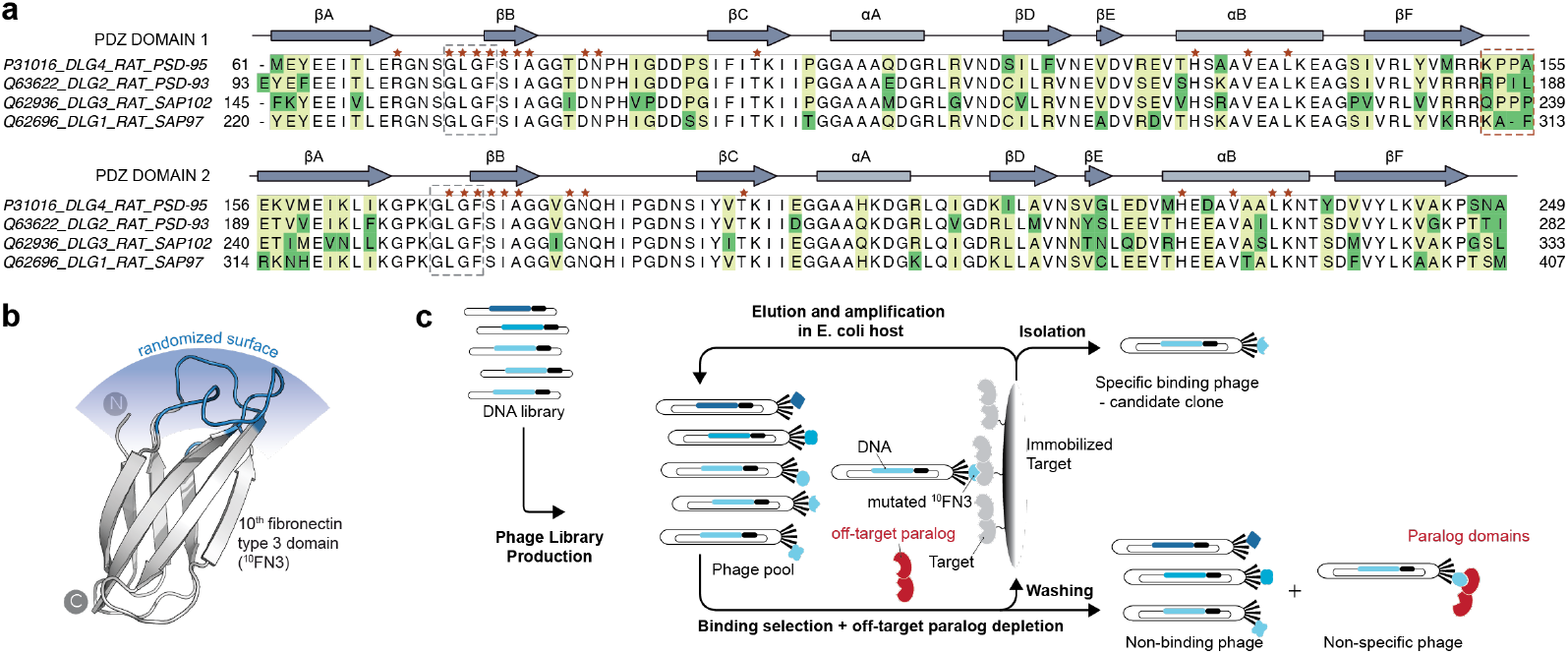
Strategy to generate PSD-95 paralog specific binders. **a**. Sequence alignment of PSD-95 paralogs tandem PDZ domains. The sequences correspond to the rat species and are provided with their associated UNIPROT entry number. The red asterisks indicate residues directly involved in the binding of partner proteins. The dark green shade highlights unique residues across the paralog family and that could be used as epitope, while the light green shade indicates residues that are present in 2 or 3 paralog proteins. **b**. Structure of the protein scaffold used for library design and selection (PDB ID 3K2M). The diversified loops (BC, DE and FG) are represented in blue. **c**. Depletion-adapted phage display strategy used for the discovery of specific PSD-95 paralog binders.

We used a phage display strategy as elaborated for the discovery of our PSD-95 specific binders, Xph15 and Xph20.(**Rimbault *et al*.,2024; Rimbault *et al*.,2019**) We started by creating two new libraries out of the robust ^10^FN3 scaffold(**Koide and Koide, 2007; Sha *et al*.,2017**) to complement the original one (**Fig. 1b**). A first set of modifications consisted in generating a library similar to our initial design(**Rimbault *et al*.,2019**) but with the mutation of a serine into a lysine group in the DE loop, to modify the charge on the diversified surface, and as well as stabilizing the S65K mutation(**Grebien *et al*.,2011**) on the opposite side of the paratope as it revealed beneficial for solubility and stability of the binders.(**Rimbault *et al*.,2019**) A second set of modifications consisted in restricting the length of the BC and FG loops diversification to the most frequently found length-combinations. After analysis of the most common loop lengths isolated from the initial library, we decided to restrict the BC length to overall 6-7 residues whereas the FG loop was kept long with 12-13 residues. The two new libraries were obtained as previously with the pFunkel method.(**Firnberg and Ostermeier, 2012**)

We used both libraries in selection campaigns performed as previously against the four tandem PDZ domains targets (PSD-95 was used here to compare libraries to the initial one). However, as paralog specificity was not obtained for target other than PSD-95 as observed initially, we modified the initial basic strictly affinity-based selection strategy to put selection pressure on specificity. This was achieved by performing a systematic depletion for the “off-target” paralogs. Practically, during each selection round the phage pools was incubated with its target in presence of micromolar concentration of the other non-targeted paralog tandem PDZ domains in a non-biotinylated form to actively remove any non-specific binders (**Fig. 1c**).

### Primary candidate evaluation

The different selections campaigns yielded distinct results (**Fig. 2a**,**b**). First, standard selections (no depletion) against PSD-95 tandem domains provided us with clones comparable to the one previously isolated when compared to Xph20 in phage-ELISA assays. This demonstrated that the new libraries are comparable in terms of quality and diversity at least from a functional point of view. As some of the new clones isolated against PSD-95 stood out with phage-ELISA responses stronger than Xph20, we kept the one with the highest response, *95*Xph142, for further characterization. In parallel, standard selection protocols failed to isolate specific binders for targets other than PSD-95, suggesting a strong “immunogenicity” from a hop-spot with unique residues for this paralog, a property not found in the others. For instance, out the clones with the strongest phage-ELISA response, a clone equally binding all paralogs was isolated from a selection against SAP102, *102*Xph68. Similarly, standard selections against PSD-93, often yielded clones with a dual PSD-93/PSD-95 specificity, *93*Xph64 and *93*Xph223 from the two library formats.

**Fig. 2.**
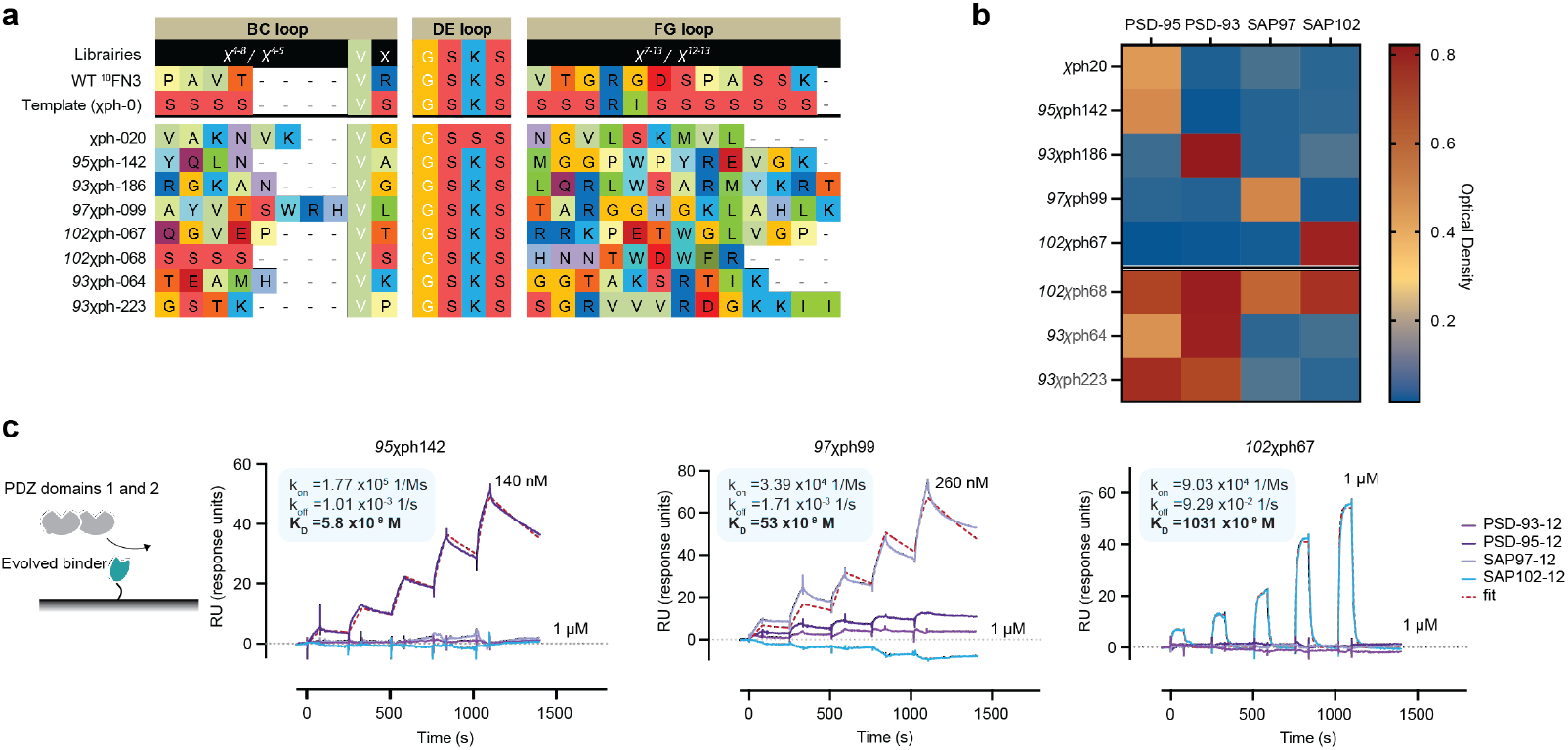
Isolation and specificity of PSD-95 paralog binders. **a**. Library design and loop sequences of the isolated clones (in the format *Target*Xph#) against paralogs first two PDZ domains. **b**. Phage-ELISA of isolated clones against tandem PDZ domains from PSD-95 family. OD, optical density at 450 nm. **c**. SPR sensorgams obtained by single-cycle kinetics of tandem PDZ domains (analyte) against immobilized biotinylated evolved binders (ligand). The reported concentrations represent the highest concentration used on the final analyte injection; the previous four injections are a series of two-fold dilutions. The colored curves represent measured data points and black lines represent the global fit obtained with a 1:1 binding model used for analysis. Kinetics values are the average of three independent titrations.

We therefore resorted to a depletion strategy (**Fig. 1c**) to focus the selection pressure on specificity properties. In practice, the addition of a freely soluble PSD-95 tandem PDZ domains construct when exposing the phage pools to their target was sufficient to allow the isolation of specific clones for the other paralogs as judged by phage-ELISA. The best responding clones for the paralogs tandem PDZ domains are presented here, for PSD-93, *93*Xph186, for SAP102, *102*Xph67 and for SAP97, *97*Xph99. We note that, while depletion with PSD-95 allowed us to isolate specific clones, it also overall significantly reduced the strength and the number of different isolated clones. As a result, except for SAP97, for which other similar clones were isolated, the two clones isolated against PSD-93 and SAP012 were by far the specific clones with the strongest response for their target. These specific clones against PSD-93, SAP102 and SAP97, together with the strong binding 95Xph142, were then further characterized.

We first focused on the determination of each clone respective epitope. As the targets are composed of two PDZ domain in tandem, phage-ELISA was first used to identify the precise domain(s) involved in the clones binding (**Supp Fig. 1**). The four specific clones (*95*Xph142, *93*Xph186, *97*Xph99 and *102*Xph67) all presented epitopes on the first PDZ domain of their respective targeted paralog. Further, the epitope of *95*Xph142 was very similar to the one of Xph20 as its binding was abolished in Xph20 presence (**Supp Fig. 1**). In parallel, *95*Xph142 binding to PSD-95 was preserved in the presence of pan-MAGUK binding *102*Xph68, confirming the idea that specific and non-specific epitopes are located on different domain regions.

The four binders were then produced in bacteria for biophysical and structural characterization. All clones, except *93*Xph186, were easily isolated. Indeed, in contrast to the others, the clone against PSD-93 could only be produced in the presence of its target and was highly unstable after the target removal. The binding properties of the other clones were assayed by surface plasmon resonance (SPR) against all tandem PDZ domains (**Fig. 2c**). Specificity of each binder was confirmed by SPR and kinetics analysis revealed fast exchanging micromolar binder for SAP102 and more stably binding, low nanomolar binders for SAP97 and PSD-95.

In solution NMR investigations of *102*Xph67 and *97*Xph99 first confirmed that in both cases PDZ domain 1 was the target. They also allowed to define molecular models for the binders-PDZ domain complexes and precise the epitopes. In both cases, epitopes were in a region distant from the PDZ domain 1 binding groove (**Supp Fig. 2**). These features indicate that the clones can be used as intrabodies to label their respective target without interfering with its PPI function.

### Affinity maturation

As the affinity of the SAP102 binder, *102*Xph67, was relatively low -in the micromolar range-and in the absence of other candidate with comparable or better binding properties as judged by phage-ELISA, we turned to an affinity maturation approach(**Thie *et al*.,2009**) to improve the clone binding properties. Random mutagenesis was performed on the whole sequence of the clone by combining the use of mutagenic dNTP analogs and error-prone PCR. Conditions were set to achieve a mutagenesis rate of about 3%. The randomized genes were incorporated into a variant of our phagemid with a pIII truncation (loss of N1 and N2). This truncation was generated in subdetection proportions in our early affinity maturation libraries and conferred a significant non-affinity-related advantage to the clones that were associated with it. In order to avoid “false” positive responses, we therefore included the truncation in our phagemid design for affinity maturation step. We suspect that this truncation already reported(**Huovinen *et al*.,2014; Larocca *et al*.,2001; Rader, 2024**) provides an advantage for phage particle production and/or improved pIII-gene fusion *vs* wild-type pIII display rate.

The randomized *102*Xph67 library was used to perform a standard phage display selection against SAP102 tandem PDZ domain with an additional off-rate selection step(**Ylera *et al*.,2013; Zahnd *et al*.,2010**) to favor recovery of binders with improved kinetic properties. Phage-ELISA and sequence analysis revealed that for best two clones (*102*Xph283 and *102*Xph285) mutations were located in the paratope (initially randomized loops that make contact with the target epitope) with an enrichment for a valine to isoleucine mutation in the FG loop (**Fig. 3a**). These mutations did not compromise the initial binder specificity as evaluated by phage-ELISA (**Fig. 3b**,**d**) and SPR. Overall, the mutations resulted in the improvement of both association and dissociation rate constants (**Fig. 3c**,**d**) and lead to a three-fold affinity improvement of the parent clone affinity.

**Fig. 3.**
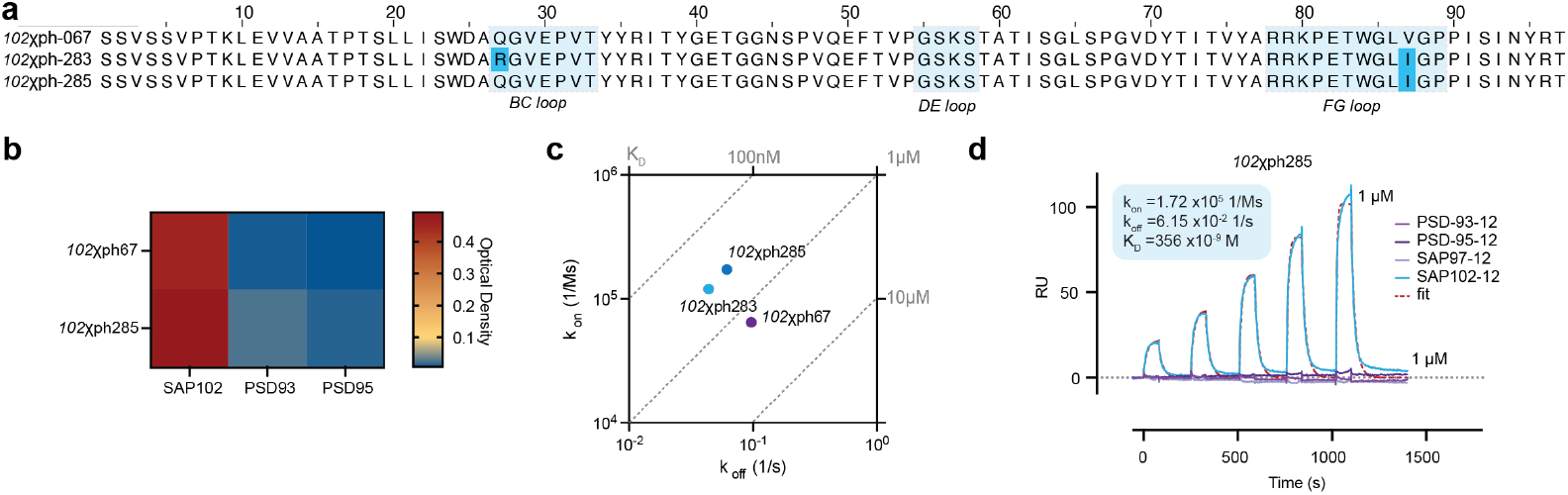
Affinity maturation of SAP102 specific binder. **a**. Sequences of the parent SAP102 evolved binder *102*Xph67 and of two isolated clones with the strongest ELISA response. Light blue shade indicates the initially diversified loops (BC, DE and FG), whereas the darker blue shade highlights the mutations. **b**. Phage-ELISA of the affinitiy matured SAP102 clone against tandem PDZ domains from PSD-95 family. OD, optical density at 450 nm. **c**. Rate Plane with Isoaffinity Diagonals (RaPID) plot for SPR data of SAP102 clones (*102*Xph67, *102*Xph283 and *102*Xph285). **d**. SPR sensorgams obtained by single-cycle kinetics of tandem PDZ domains (analyte) against immobilized biotinylated affinity matured SAP102 binder (ligand). The reported concentrations represent the highest concentration used on the final analyte injection; the previous four injections are a series of two-fold dilutions. The colored curves represent measured data points and black lines represent the global fit obtained with a 1:1 binding model used for analysis. Kinetics values are the average of three independent titrations.

### Cellular target recognition

We next sought to confirm the evolved binders capacity to recognize and engage with their respective target when those targets are expressed as full-length proteins in cells. Indeed, as the selection was performed on a fragment of the protein (albeit a supramodule fragment, which according to the current structural understanding operate relatively independently to the other domains(**Feng and Zhang, 2009; Laursen *et al*.,2022; Laursen *et al*.,2020**)), there remains a possibility that the epitope is occluded in the intact protein, either by intra- or inter-molecular interactions or by post-translational modifications. We also considered that the binders themselves might be unstable and therefore nonfunctional in the cellular environment.

We used a FRET/FLIM (Förster resonance energy transfer/fluorescence-lifetime imaging microscopy) assay to report on the interactions between the binder and its target. The FRET system was based on one previously developed to investigate divalent ligands(**Sainlos *et al*.,2011**) and PSD-95 binders(**Rimbault *et al*.,2019**). The donor fluorescent protein, eGFP, was inserted after the second PDZ domain in PSD-95, PSD-93, SAP102 or SAP97 (**Fig. 4a**). The acceptor, mCherry, replaces the C-terminal PDZ domain-binding motif of the transmembrane protein Stargazin, and is followed by a 20-amino-acid linker and the evolved binder. Proximity of the end of the second PDZ domain to the first PDZ domain, which is the targeted module for all specific binders, should allow to detect interactions by modulation of the donor lifetime as observed previously for PSD-95 binders(**Rimbault *et al*.,2019**). Non-modified Stargazin (intact PDZ domain-binding motif) and Xph0, a naïve non-binding clone, were used as positive and negative control respectively. All constructs were expressed in COS-7 cells and FRET accessed by measurement of the donor lifetime. For PSD-95, the donor lifetime was significantly shortened with Xph20, *95*Xph142 and *93*Xph64 as expected from initial selectivity characterization. The stronger binding affinity of *95*Xph142 over Xph20 were confirmed by the stronger FRET effect of the new clone. No detectable interaction was detected for *93*Xph186 or *97*Xph99. Similar results were obtained for PSD-93 and SAP102, with only the expected binders, *93*Xph186 and *93*Xph64 for PSD-93 and *102*Xph67 for SAP102, modifying the donor lifetime. These results beside confirming the cellular specificity of the clones also demonstrated that despite its instability in bacteria, *93*Xph186, can be efficiently produced in mammalian cells and be functional. Finally, no binding was observed for the clone against SAP97 (other clones were tested but yielded the same results) suggesting that the epitope recognized by SAP97 specific clones is not accessible in cells with the full-length protein. Overall, except for SAP97 binders, these results confirmed the capacity of the evolved binders to recognize and bind specifically their target in cells.

**Fig. 4.**
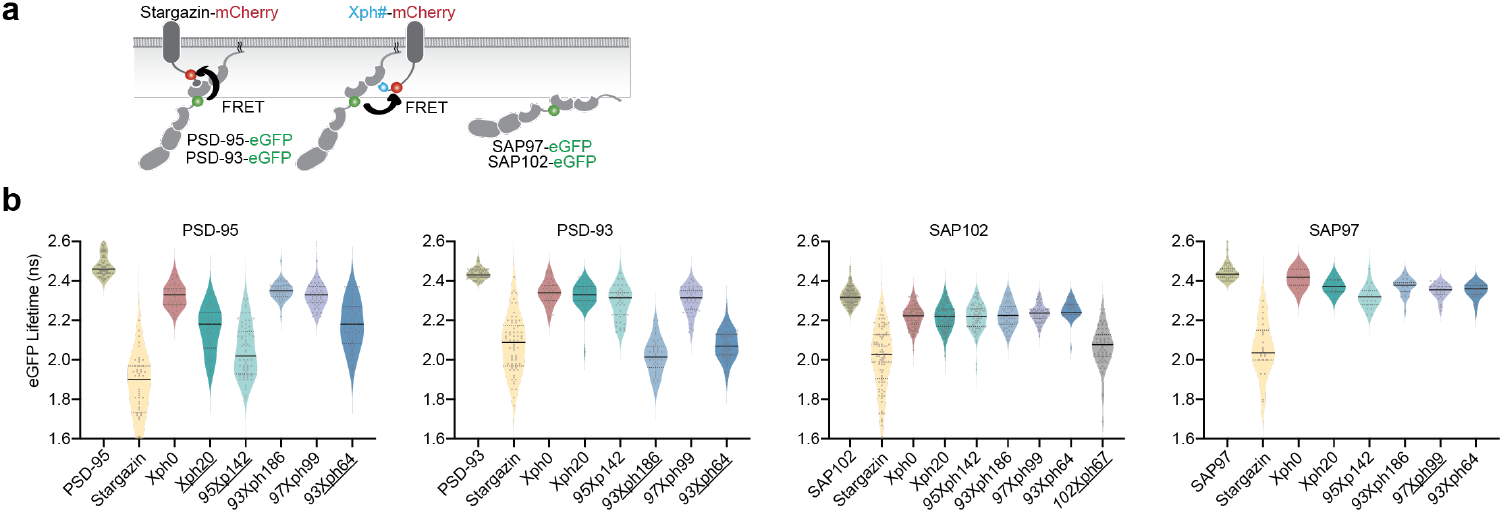
Evolved binder full-length target recognition in a cellular environment. **a**. Schematic of the FRET systems used for measurement of the donor lifetime (FLIM). **b**. Lifetime of eGFP inserted in PSD-95, PSD-93, SAP102 and SAP97 in presence of the indicated acceptor-containing protein constructs. Stargazin(-mCherry) as a positive control and membrane anchored binders with Xph0 as negative control, Xph20 as reference. Binders from which a interaction with the target is expected are underlined. Violon plots show median, first and third quartile, with whiskers extending to the minimum and maximum and all individual data points (each corresponding to a single cell) pooled from at least three independent experiments.

### Intrabody imaging probes for PSD-95 paralogs

After selection and characterization of binders across the PSD-95 paralog family, we designated *95*Xph142, *93*Xph186 and *102*Xph285 as lead candidates to develop new imaging probes to label endogenous PSD-95, PSD-93 and SAP102, respectively. Indeed, we showed that they were efficiently and specifically binding their respective target in cells and that their epitopes (known for *95*Xph142 and *102*Xph285; partially inferred for *93*Xph186) are consistent with absence of impact on the PPI module activity of the MAGUK proteins. These features make them well ideally suited for the development gene-encoded intrabodies for advanced imaging. While no such tools exist for PSD-93 and SAP102, the interest for a new PSD-95 binders reside here in complementing the already existing toolset (Xph15 and Xph20) with a binder with different kinetic properties(**Rimbault *et al*.,2024**).

The three binders were therefore cloned into plasmids with a regulated expression systems as previously used for PSD-95 probes.(**Gross *et al*.,2013; Rimbault *et al*.,2024**) Probe expression is hence regulated by fusing a transcriptional repressor to a zinc finger and placing the cognate zinc-finger binding motif upstream of the reporter gene in the expression plasmid. In this setup, the fluorescent protein (FP) monitors the interaction between the binding module and its target, while the regulation module curbs recombinant binder levels to prevent overexpression relative to the endogenous target. The three intrabodies were expressed in rat neuron primary hippocampal cultures and observed after 15 days in vitro (DIV 15). All intrabodies expressed similarly, with no sign of toxicity. They all showed characteristic synapse enriched-structures (**Supp Fig. 3**) as expected for synaptic scaffold proteins binders.

Their labeling properties were compared to standard antibody staining (**Fig. 5**). Both for *102*Xph285 (**Fig. 5a-c**) and for *93*Xph186 (**Fig. 5d-f**) the intrabody signal visually correlated with the labeling obtained with specific antibodies. Line scan intensities for the two channels confirmed this observation. To complement the evaluation, we generated guide RNA to knock down SAP102 (both gRNA target regions upstream of the first PDZ domain) and confirm SAP102 intrabody specificity. The two SAP102 guide RNAs or a control guide RNA with a TdTomato reporter and hSpCas9 enzyme were nucleofected into cortex culture together with the regulated *102*Xph285-eGFP. At DIV 20, cells fixed and labelled with an antiSAP102 antibody. Comparison of the two conditions clearly demonstrates first the SAP102 knock out (**Fig. 6**) with clear loss of the antibody signal for the two gRNAs expressing cell and not for the control gRNA. In addition, this loss of signal is mirrored by the SAP102 probe, *102*Xph285, with a signal above auto-fluorescence contained in the nucleus as expected from the regulation system, clearly proving its specific binding properties.

**Fig. 5.**
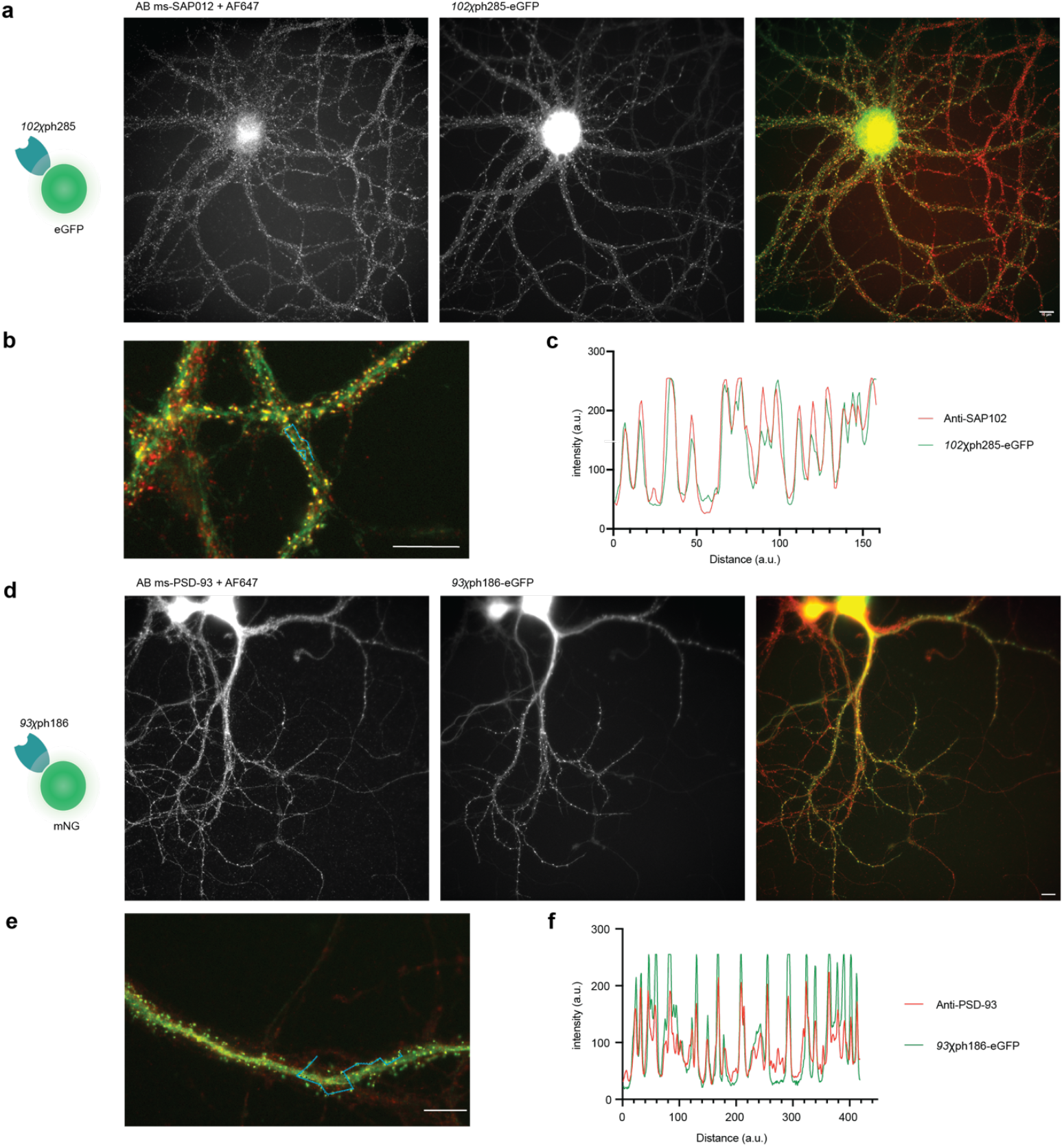
Evaluation of evolved binders as intrabody fluorescent reporter probes. **a**. Representative epifluorescence images of the eGFP-fused *102*Xph285 binding module *vs* immunostaining of endogenous SAP102 in rat hippocampal primary cultures (scale bar 10 µm). **b**. and **c**. Line scan measurement of antibody (red) and intrabody (green) labeling. The line scan is represented in **b**. with a blue line and corresponding intensities plotted in **c**. (scale bar 10 µm). **d**. Representative epifluorescence images of the mNeonGreen-fused *93*Xph186 binding modules *vs* immunostaining of endogenous PSD-93 in rat hippocampal primary cultures (scale bar 10 µm). **e**. and **f**. Line scan measurement of antibody (red) and intrabody (green) labeling. The line scan is represented in **e**. with a blue line and corresponding intensities plotted in **f**. (scale bar 10 µm).

**Fig. 6.**
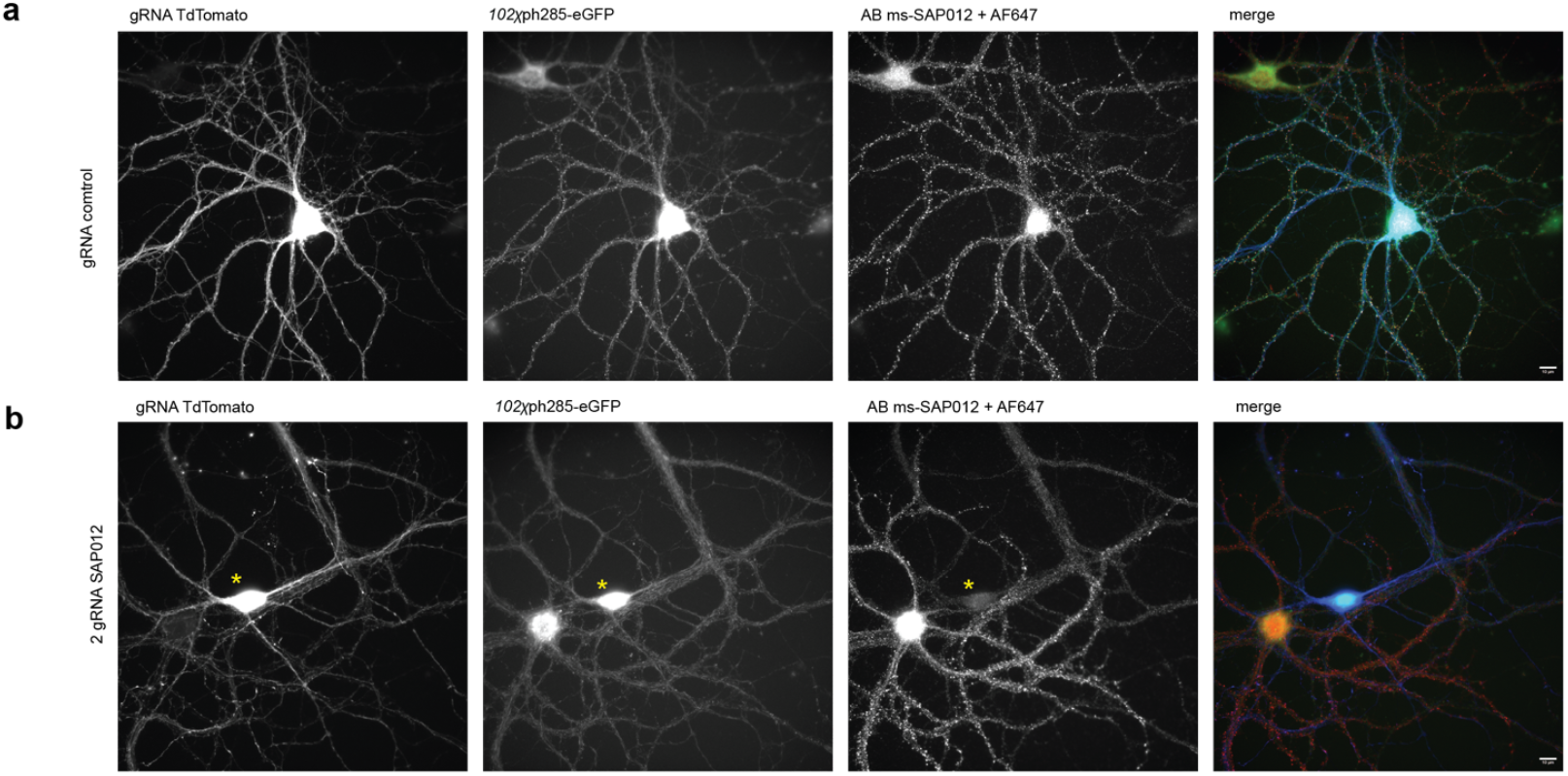
Evaluation of *102*Xph285 intrabody specificity. **a**. and **b**. Representative epifluorescence images of the FP-fused *102*Xph285 *vs* immunostaining of endogenous SAP102 in presence of CRISPR-Cas9 and either control guide RNA (**a**.) or 2 guide RNA FP against SAP102 (**b**.) each with TdTomato reporter in rat cortex primary cultures (scale bar 10 µm). The yellow asterisk indicates the guide RNA and intrabody expressing neuron. The neighboring neuron indicate the level of green autofluorescence.

## Discussion

Our selections demonstrate that paralog-specific recognition within a highly conserved PDZ domain supramodule is achievable when selection pressure explicitly favors specificity. Introducing soluble “off-target” paralogs during panning enriched clones that discriminate PSD-93 and SAP102, while standard affinity-only selections predominantly recovered PSD-95-biased or pan-MAGUK binders. Mechanistically, all specific clones mapped to epitopes on PDZ1, consistent with the availability of paralog-unique surface residues.

We converted the lead binders—*95*Xph142 (PSD-95), *93*Xph186 (PSD-93) and *102*Xph285 (SAP102; further optimized by affinity maturation)—into regulated intrabody probes. In neurons, these probes reproduced synaptic labeling seen with antibodies, and SAP102 probe specificity was further confirmed by CRISPR knock-down. The zinc-finger–based regulation(**Gross *et al*.,2013**) kept unbound probe levels low, minimizing overexpression artifacts and echoing our earlier PSD-95 work.(**Rimbault *et al*.,2024**) Together, these results establish a practical pipeline from phage selection to gene-encoded imaging tools for multiple MAGUK paralogs.

Our findings also align with prior principles derived for PSD-95: targeting non-functional surfaces away from the PDZ domain binding groove favors minimal perturbation, and specificity can be anchored by a small set of divergent residues (*e*.*g*., F119 in PSD-95).(**Rimbault *et al*.,2019**) In PSD-95, Xph binders neither altered AMPAR currents nor receptor mobility, and regulated expression tightly coupled intrabody levels to the endogenous target—design features we now extend to other paralogs.

A notable limitation emerged with SAP97: despite selection of PDZ1-directed clones, no interaction was detected with the full-length protein in cells, suggesting epitope occlusion and highlighting the importance of conformational context. Selecting against the entire PDZ1–2 supramodule was originally conceived to avoid such issues, yet higher-order interactions or post-translational modifications in full-length proteins can still mask accessible surfaces. Future selections that combine depletion with alternative scaffolds or epitope re-targeting, may overcome this barrier.

Looking ahead, these paralog-specific intrabodies enable multiplexed, low-perturbation interrogation of MAGUK family members in living neurons. Orthogonal regulation systems are available for combinatorial deployment, and the binders can be repurposed to target sensors or enzymatic modules (*e*.*g*., SNAP-tag, GCaMP) to defined subsynaptic locales. This toolkit should facilitate tests of paralog-specific roles in nanoscale synapse organization—including trans-synaptic nanocolumn alignment—and help relate composition to function across synapse subtypes.

Overall, by combining specificity-focused selection with regulated expression, we expand endogenous, gene-encoded imaging from PSD-95 to its paralogs, providing a coherent framework for dissecting paralog diversity at the synapse with minimal perturbation.

## Materials and Methods

### Plasmid construction

The plasmids generated and the primers used in this study are listed in **Supplementary Tables 1** and **2**, respectively. Plasmids for protein production were previously described (**Rimbault *et al*.,2019; Sainlos *et al*.,2009**). Briefly, for bacterial expression, the first two PDZ domains of PSD-95 were subcloned into pET-NO to produce a N-terminal fusion with an octa-His tag and a TEV protease cleavage site. The Xph clones were subcloned into the pIGc vector to generate C-terminal fusions with a deca-His tag. For FRET experiments, PSD-95- and paralogs-eGFP fusions and stargazin-mCherry were previously described (**Rimbault *et al*.,2019; Sainlos *et al*.,2011**). TM-mCherry-Xph constructs were obtained by digesting and ligating the corresponding Xph from pET-IG (resulting in a N-terminal linker of 19 amino acids) into Stargazin-mCherry plasmid C-terminally to mCherry (replacement of Stargazin C-terminus) between NdeI and XhoI sites.

For imaging, Xph binders were subcloned into pCAG_PSD95.FingR-eGFP-CCR5TC (gift from Don Arnold, USC, Addgene #46295) (**Gross *et al*.,2013**) using KpnI and BglII restriction sites. Other fluorescent modules, mRuby2 (gift from Michael Lin, Addgene #40260) (**Lam *et al*.,2012**), mNeonGreen (obtained by gene synthesis, Eurofins) (**Shaner *et al*.,2013**), HaloTag (Promega, cat no G7971) and SNAPf (New England Biolabs, cat no N9183S) were next inserted in place of eGFP in the corresponding vector using BglII and NheI sites after an initial modification of the source vectors to introduce an NheI site between the fluorescent module and CCR5 ZF. AAV expressing vector containing Xph binders were subcloned into AAV-Syn-PSD95.FingR-eGFP-CCR5TC (gift from Xue Han, Addgene plasmid # 125693) (**Bensussen *et al*.,2020**) by replacing the PSD95.FingR coding sequence using the NheI and SphI restriction sites.

The CRISPR target sequences were all 20-nucleotide long and followed by a protospacer adjacent motif (PAM). The guide RNA sequence to Knock-down SAP102 expression were designed in silico using the CRISPOR design tool software (https://crispor.gi.ucsc.edu/). We chose to target the exon1, near ATG and the exon 5 coding for PDZ1 domain of the SAP102 rat sequence (Q62936) (gene Dlg3 Gene ID: 58948) to remove the N-terminal part up to the PDZ1.Two guide RNA (gRNA) sequence were selected, respective sequences 5’-CGAGTGCTATGAGGTGACCC -3’ and 5’-TGAGGTGGACGTGTCCGAGG-3’. As a control we used a sequence from a gecko bank: 5’-ATATTTCGGCAGTTGCAGCA-3’. gRNAs were cloned into the vector pSpCas9(BB)–2A-GFP (PX458) (Addgene cat#48138). The GFP sequence was replaced by TdTomato sequence using the HD-In-Fusion kit (Takarabio) with primers 5’ TGACGTCGAGGAGAATCCTGGCCCAGTGAGCAAGGGCGaggagg3 and ‘5’ GAGGCTGATCAGCGAGCTCTAGttaCTTgtacAGCTCGTCCATgcc 3’ to amplify TdTomato and 5’ TAACTAGAGCTCGCTGATCAG 3’ and 5’ TGGGCCAGGATTCTCCTCG 3’ to linearized the vector. The gRNA sequence was annealing and cloned in BbsI restriction sites following the Zhang’s Lab protocols (**Cong *et al*.,2013**). Vector pSpCas9(BB)-2A-GFP (PX458) was a gift from Feng Zhang (Addgene plasmid # 48138; http://n2t.net/addgene:48138; RRID:Addgene_48138).(**Ran *et al*.,2013**)

### Library construction

Libaries were constructed as previously described(**Rimbault *et al*.,2019**). The ssDNA was obtained for E. coli CJ236 dut-ung-strain (TaKaRa, E4141S). Combinatorial libraries in which the BC and FG loops of 10FN3 were diversified were constructed by using pFunkel mutagenesis(**Firnberg and Ostermeier, 2012**), inspired by Kunkel mutagenesis.

#### Error-prone PCR library generation

Random mutagenesis was carried out in a 50 μl reaction with the JBS dNTP-Mutagenesis Kit (Jena Bioscience) with 0.5 mM of each dNTP, 1.1 mM of each natural dNTPs, 0.5 μM primers and 25 fmol of DNA template. The PCR reaction was run with the following settings: (1) 1min at 92 °C, (2) 1.5min at 55 °C, (3) 5min at 72 °C, (4) 30x to step 1. A second PCR reaction was performed with 1 ul of the PCR product from the first PCR reaction, 0.5 μM primers and 0.5 mM each dNTP. In parallel, vector preparation was performed by PCR. In a 50 μl reaction, 1 ng of vector DNA was combined with 0.5 μM primers, 0.2 mM of each dNTP, 0.02 U/μl of Q5U® Hot Start High-Fidelity DNA Polymerase (New England Biolabs), 1× Q5U® buffer, and X μM of DMSO. The PCR reaction was run with the following settings: (1) 1min at 98 °C, (2) 20sec at 98 °C, (3) 20sec at 60 °C, (4) 4min at 7 2°C, (5) 30x to step 2, (6) 5min at 72 °C. Both PCR products were treated with DpnI for 1 hour at 37 °C, then purified using the GeneJET PCR Purification Kit (ThermoScientific).

The NEBuilder® HiFi DNA Assembly reaction was then performed with 0.065 pmol of linearized vector and 0.13 pmol of the diversified insert (from random mutagenesis), according to the manufacturer’s instructions, for 15 min at 50 °C. The assembled product was desalted and concentrated using Amicon Ultra-0.5 Centrifugal Filter Devices with a 30K molecular weight cut-off (Merck Millipore), followed by lyophilisation and resuspension in water to a final concentration of 10 ng/μl for subsequent electroporation into TG1 electro competent cells (Lucigen).

In two pre-chilled 0.1 cm cuvettes (BioRad), the DNA library (10 ng) was electroporated into 25 µL of TG1 cells at 1800V with an electroporator (BioRad MicroPulser) and then incubated with 1 ml prewarmed Recovery medium (Lucigen) for 1 h at 37 °C with shaking at 250 rpm. A total of 10 µL of the electroporated cells were kept for serial dilutions and plating on 2x YT agar-plate with 50 µg mL^−1^ carbenicillin and 10 mM glucose to determine the electroporation efficiency and estimate the library size. The entire remaining volume was then plated on dishes (Greiner bio-one, 145 mm × 145 mm × 20 mm) containing 2x YT agar-plate with 50 µg mL^−1^ carbenicillin, 10 mM glucose and incubated overnight at 37 °C. Each plate was recovered with 4 ml of 2x YT medium and scraped to recover library-containing TG1 cells. Two more washes with 2 ml of 2x YT medium were made to ensure maximal library recovery. Bacteria were centrifuged at 5000 × g for 10 min at 4 °C and suspended in fresh 2x YT medium for two times. Bacteria were then flash-frozen with liquid nitrogen after the addition of 20% sterile glycerol and were kept at −80 °C for later use.

### Biopanning procedure

Libraries and panning were performed as previously described(**Rimbault *et al*.,2019**) with following some modifications. For depletion, 10 µM non-biotinylated PSD-95 tandem PDZ domains were added when phage pools were incubated with target immobilized on magnetic beads. For maturation, in order to apply selection pressure based on off-rate, beads were then incubated for 10 minutes at 37 °C in 1 mL of PBS (0.2% Tween 20) containing a competing off-rate molecule (not biotinylated SAP-102 PDZ domains 1 and 2).

#### Phage-ELISA

Phage-ELISAs were performed as previously described(**Rimbault *et al*.,2019**) with horseradish-peroxidase-conjugated anti-M13 monoclonal antibody (either Antibody design AS003 or SinoBiological 11973-MM05T-H, diluted at 1:1,1000 to 1:5,000 in PBT).

### Protein production

Proteins were expressed and purified as previously described (**Rimbault *et al*.,2019**). Briefly, His-tagged proteins were either produced in *E. coli* BL21 CodonPlus (DE3)-RIPL competent cells (Agilent, 230280) using auto-induction protocols (**Studier, 2005**) at 16 °C for 20 h or in BL21 pLysY (New England Biolabs, C3010I) for isotopically-labelled proteins with IPTG induction for 16 h at 20 °C. Proteins were first isolated by IMAC using Ni-charged resins then further purified by size exclusion chromatography (SEC). An intermediate step of affinity tag removal by incubation with the TEV protease was added before the SEC step for isotopically-labelled proteins. The recovered proteins were concentrated, aliquoted and flash-frozen with liquid nitrogen for conservation at -80 °C.

For biotinylated proteins, tandem PDZ domains and Xph clones were cloned into pbIG or pIGb vectors by inserting the desired genes between BamHI and XhoI or NdeI and XhoI restriction sites respectively. Home-made E. coli BL21 (DE3) competent cells (source ThermoFisher Scientific, C600003) containing the pACYC-mCh-BirA plasmid were transformed with the gene of interest into pbIG or pIGb. Transformed bacteria were selected on LB plate containing chloramphenicol and kanamycin for maintaining the two expression plasmids. 5 mL of LB starter cultures with appropriate antibiotics were inoculated into 300 mL of ZYM-5052 auto-inducing media containing the appropriate antibiotics (30 µg.ml^-1^ for kanamycin and 34.5 µg.ml^-1^for chloramphenicol). Cultures were grown in sterile glass vessel 5 h at 37 °C at 250 rpm. Biotin (50 µM) was then added to the medium and the temperature was lowered to 16 °C for expression of the target proteins (∼20 h). Proteins were purified as described for His-tagged proteins. Biotinylation yields were evaluated using either the Pierce Biotin quantitation Kit following the manufacturer protocol (ThermoFisher Scientific) or by a SDS-PAGE assay(**Sorenson *et al*.,2015**).

### Surface Plasmon Resonance

Surface plasmon resonance measurements were performed on a BIAcore X100 instrument with analysis temperature set to 25 °C. CAP Sensor chips (GE Healthcare) that allow the reversible capture of biotinylated ligands as an immobilization system were used. During the experiments, reagents were kept at room temperature. Immobilization levels were optimized to reflect a compromise between minimal surface density of the ligand to avoid rebinding effects and generation of exploitable sensorgrams when possible.

Sensorgrams were collected using single cycle kinetics as tandem PDZ domains were injected at various concentrations (using two-fold dilutions unless otherwise stated) in PBS containing 500 mM NaCl (pH 7.4), supplemented with 0.1 % Tween-20 at a flow rate of 30 µL.min^-1^. All sensorgams were double referenced prior to analysis by first subtracting data from a reference flow cell functionalized with the capture reagent but on which no ligand was attached and then subtracting a blank cycle where buffer was injected instead of the protein sample. The kinetic data were analyzed using a 1:1 Langmuir binding model of the BIAevaluation software with the bulk refraction index (RI) kept constant and equal to 0 and the mass transfer constant (tc) kept constant and equal to 10^8^.

### NMR spectroscopy

NMR spectra were recorded at 298 K using a Bruker Avance III 700 MHz spectrometer equipped with a triple resonance gradient standard probe. Topspin version 4.1 (Bruker BioSpin) was used for data collection. Spectra processing used NMRPipe (**Delaglio *et al*.,1995**). with analysis by using Sparky 3 (T. D. Goddard and D. G. Kneller, University of California).

### FRET/FLIM assays

FRET/FLIM assays were performed as previously described (**Rimbault *et al*.,2019**). Briefly, COS-7 cells (ECACC-87021302) in DMEM medium supplemented with Glutamax and 10 % FBS were transfected using a 2:1 ratio X-treme GENE HP DNA transfection reagent (Roche) per µg of plasmid DNA with a total of 0.5 µg DNA per well. Experiments were performed after 24 h of expression. Coverslips were transferred into a ludin chamber filled with 1 ml fresh Tyrode’s buffer (20 mM Glucose, 20 mM HEPES, 120 mM NaCl, 3.5 mM KCl, 2 mM MgCl_2_, 2 mM CaCl_2_, pH 7.4, osmolarity around 300 mOsm.kg^-1^and pre-equilibrated in a CO_2_ incubator at 37 °C).

Experiments with full-length PSD-95 paralogs were performed using the frequency domain analysis (LIFA) method a Leica DMI6000 (Leica Microsystems, Wetzlar, Germany) equipped with a confocal Scanner Unit CSU-X1 (Yokogawa Electric Corporation, Tokyo, Japan). The FLIM measurements were done with the LIFA fluorescence lifetime attachment (Lambert Instrument, Roden, Netherlands), and images were analyzed with the manufacturer’s software LI-FLIM software.

Lifetimes were referenced to a 1 µM solution of fluorescein in Tris-HCl (pH 10) or a solution of erythrosin B (1 mg.ml^-1^) that was set at 4.00 ns lifetime (for fluorescein) or 0.086 ns (for erythrosin B). For competition experiments, only cells presenting a high level of expression of the competitor or control as measured by mIRFP670 fluorescence were taken into consideration.

### Rat hippocampal cultures

All experiments were performed on E18 rat dissociated hippocampal banker culture. Sprague-Dawley rats (Janvier Labs) of either sex were euthanized in accordance with the European 2010/63/EU directive, Annex IV Methods of killing animals. All experiments were performed in accordance with the European guidelines for the care and use of laboratory animals, and the guidelines issued by the University of Bordeaux animal experimental committee (CE50; Animal facilities authorizations A3306940, A33063941). Dissociated hippocampal neurons from E18 embryos were prepared as described previously(**Kaech and Banker, 2006**) at a density of 200,000 cells per 60 mm dish on poly-L-lysine pre-coated coverslips and which are put after 2h in the ‘sandwich configuration’ in dishes containing confluent astrocytes. Cultures were maintained at 36.5 °C, 5% CO_2_ in Neurobasal™ Plus medium (Thermofisher Scientific cat # A3582901) supplemented with 2 mM L-glutamine and B-27™ Plus Neuronal Culture System (Thermofisher Scientific cat # A3582801), during at least 3 days. After 3 days, media was supplemented with cytosine arabinoside “Ara-C” (5 μM) to inhibit glial cell proliferation. The cultures were either (i) left in this medium, renewed by a third twice a week.

### Gene delivery

For SAP102 immunostaining, experiments, 500K of rat hippocampal neurons from E18 embryos were nucleofected at DIV 0 before plating with 2 μg of Xph285 DNA per 60 mm dish containing four 18mm glass using Nucleofector system (Lonza) in 100 μl Single Nucleocuvette with P3 Primary Cell 4D-Nucleofector X Kit and HV hippocampal neuron program.

For SAP102 CRISPR/Cas9 knock-down experiments, 1µg of each plasmid carrying a gRNA were co-nuclefected with the Xph285 DNA plasmid.

For PSD-93 immunostaining experiments, rat hippocampal neurons from E18 embryos were transfected with 4 µg of Xph186 DNA per 60 mm dish containing four 18mm glass coverslips using a standard calcium phosphate protocol at 8 DIV.

### Immunostaining

At 16–20 DIV, E18 rat Banker cultures expressing individual eGFP-tagged Xph or were stained either with a with mouse monoclonal anti-SAP102 (antibodiesinc /Neuromab Cat# 75-058) at 2 μg/ml final concentration or mouse monoclonal anti-PSD-93 (antibodiesinc /Neuromab Cat# 75-057) at 10 μg/ml final concentration. Briefly, neurons on coverslips were fixed 10 min using PFA 4%, washed with PBS, permeabilized with PBS-Triton-0.1% during 5 min, and washed again. After blocking with PBS-BSA 1%, neurons were stained with the primary antibody and after three washes with the respective secondary antibody Anti-mouse Alexa 647 (goat polyclonal Thermo Scientific Cat# A21235) or Anti-mouse Alexa 568 (Donkey polyclonal Thermo Scientific Cat# A10037) for 45 min each. Neuron coverslips were mounted with Fluoromont G™mounting medium reagent (Thermo Scientific, Cat# 00-4958-02). Images were acquired on a Leica DM5000 (Leica Microsystems) with a HCX PL APO.63 oil NA 1.40 objective, a LED SOLA Light (Lumencor, Beaverton) as fluorescence excitation source and a Flash4.0 V2 camera (Hamamatsu Photonics, Massy, France).

## Acknowledgments

This research was financially supported by grants from the Centre National de la Recherche Scientifique, the Conseil Régional d’Aquitaine, the France BioImaging national infrastructure (grant ANR-10-INBS-04), the Agence Nationale de la Recherche (OptoXL, ANR-16-CE16-0026) (AFFLIGEM, ANR-21-CE44-0013) to C.P, CM and M.S, the European Research Council to D.C. We also thank the IINS cell culture facility for technical assistance, the Biochemistry and Biophysics Core Facility of the Bordeaux Neurocampus both funded by the Labex BRAIN (ANR-10-LABX-43). We acknowledge Andrea Lopez-Castillo for help with protein production. Financial support from the IR-RMN-THC Fr3050 CNRS for conducting the research is gratefully acknowledged.

## Author contributions

M.S. designed the research and wrote the article with input from all authors. E.R. performed the selection and biophysical experiments and generated the constructs with the help of C.B., C.R. M.D. performed the affinity maturation experiments. V.T., A.E. C.M. performed the NMR experiments. C.P. performed the FRET experiments. CB performed the cellular imaging experiments with help from SD from CRISPR KO. M.S. coordinated and oversaw the research project. All authors discussed the results and commented on the manuscript.

## Declaration of Generative AI and AI-assisted technologies in the writing process

During the preparation of this work the authors used ChatGPT in order to improve clarity and succinctness. After using this tool/service, the authors reviewed and edited the content as needed and take full responsibility for the content of the publication.

## Competing interests

The authors declare no competing interest.

## Notes

### Competing Interest Statement

The authors have declared no competing interest.

